# Computing the geometry of phase–amplitude coupling: A new moment-based framework

**DOI:** 10.64898/2026.09.15.751789

**Authors:** Mahmoud Keshavarzi, Usha Goswami

## Abstract

Phase–amplitude coupling (PAC) is widely used to quantify interactions between neural oscillations, yet cannot establish whether two coupling profiles share the same geometry, phase polarity or modal structure. Here we demonstrate that commonly-used PAC metrics constitute many-to-one mappings from phase–amplitude distributions to low-dimensional summaries. This results in distinct coupling geometries yielding identical PAC values. We propose formalising PAC as a hierarchy of representations, introducing a circular-moment decomposition that preserves richer geometric information. Simulations demonstrate that distinct coupling geometries produce identical Modulation Index (MI) values. Analyses of EEG data acquired during adult language listening reveal substantial geometric variability among PAC profiles with matched MI values, validating the moment-based approach. Applied to a child-language dataset, the framework identifies differences in phase geometry of delta-low gamma PAC in children with developmental language disorder, potentially uniting competing neuroscientific theories. We propose a representation-selection framework for future PAC research.

## Introduction

A central challenge in computational neuroscience is whether summary statistics faithfully represent the structures they are intended to capture. When a complex, high-dimensional object is reduced to a scalar or low-dimensional descriptor, information is necessarily discarded. The consequences of this compression depend on whether the discarded information is irrelevant to the scientific question at hand or whether it encodes properties that are essential for mechanistic interpretation. This problem arises across statistical inference, information theory, and machine learning, where different latent states may generate identical observations and therefore cannot be distinguished by their summary representations alone (Cover and Thomas, 2006; Bishop, 2006). Whether widely used scientific measurements share this property has received comparatively little attention.

Phase–amplitude coupling (PAC), in which the amplitude of high-frequency neural activity is modulated by the phase of a slower oscillation, exemplifies this problem. PAC has been investigated across species, recording modalities, and cognitive domains and is among the most widely-used measures of multiscale neural interaction (Canolty et al., 2006; Jensen and Colgin, 2007; Siegel et al., 2009; Tort et al., 2010; Axmacher et al., 2010; Canolty and Knight, 2010; van der Meij et al., 2012; Lisman and Jensen, 2013; Szczepanski et al., 2014; Watrous et al., 2015; Lizarazu et al., 2019; Daume et al., 2024). Regarding speech processing, delta-beta and theta-gamma PAC play a central role in auditory neuroscience and neurolinguistics. Low-frequency oscillations are thought to provide a temporal scaffold for speech and language processing by organising higher-frequency activity according to the rhythmic structure of incoming speech, linking slower prosodic and syllabic timescales to faster phonemic and lexical computations (Giraud and Poeppel, 2012; Hyafil et al., 2015; Arnal et al., 2015; Dogonasheva et al., 2025). Given the central mechanistic role accorded to PAC in adult speech and language processing, PAC is frequently examined in relation to developmental language and reading difficulties. Yet surprisingly, PAC appears intact in both developmental language disorder (DLD) and developmental dyslexia, despite measurable linguistic difficulties (Power et al., 2016; Lizarazu et al., 2023; Keshavarzi et al., 2024). This appears puzzling given the central mechanistic role of PAC regarding adult speech comprehension and speech processing.

As PAC is not directly observable, it must be estimated from neural recordings using quantitative metrics. Proposed measures include the modulation index (MI, Tort et al., 2010), mean vector length (MVL, Canolty et al., 2006), phase-locking approaches (Penny et al., 2008), and generalised linear models (Kramer and Eden, 2013). Despite methodological differences, all of these approaches compress a complex phase–amplitude relationship into one or a few descriptors. Methodological work has improved the reliability of PAC estimation by addressing confounds such as filtering artefacts, waveform asymmetry, and non-stationarity (Aru et al., 2015; Jensen et al., 2016; Cole and Voytek, 2017). A more fundamental question, however, has remained largely unexamined: what information about the underlying phase–amplitude structure is retained or discarded when a metric is computed? In representational terms, this is a question of identifiability, namely whether the mapping from phase–amplitude distributions to metric values is one-to-one or whether multiple distinct coupling structures can produce the same output.

An initial indication that information loss in PAC metrics may be substantial and consequential was recently established by Keshavarzi (2026). Combining mathematical analysis with empirical EEG data, it was shown that MI-based PAC is invariant to 180° inversion of the low-frequency phase. Accordingly, temporally-opposite coupling structures, in which high-frequency amplitude is maximal at opposite phases of the slow oscillation, can yield numerically identical MI values. If significant phase-difference is lost during MI computation, this could explain the puzzling data regarding intact PAC in developmental disorders of language. Keshavarzi (2026) establishes a concrete case in which a widely-used PAC metric discards information that is directly relevant to mechanistic interpretation.

The present study asks whether this limitation reflects a general representational characteristic of PAC metrics. The phase–amplitude relationship is formalised as a probability distribution over low-frequency phase, and PAC measures are treated as mappings from this distribution to lower-dimensional representations. Within this formulation, scalar measures such as MI are shown to be many-to-one mappings. This demonstrates that distinct phase–amplitude geometries, including unimodal, bimodal, and multimodal coupling structures, can yield identical metric values despite representing different temporal organisations. Phase-inversion invariance therefore emerges as one example of a broader class of non-identifiabilities arising when complex phase–amplitude distributions are compressed into scalar summaries. If MI cannot distinguish between opposite temporal organisations of cross-frequency coupling, MI values alone cannot support inferences about whether the phase alignment of neural activity with rhythmic input is intact in language disorders.

To quantify the information preserved by alternative representations, we introduce a circular-moment framework in which the phase–amplitude distribution is described by its Fourier coefficients. This formulation establishes a hierarchy of representations between maximally-compressed scalar summaries and the complete phase–amplitude distribution. Conventional PAC metrics emerge as low-dimensional projections of this richer space, whereas successive moments capture increasingly detailed geometric information about the underlying coupling structure. In the limit of the complete moment set, the original distribution is recovered exactly. The framework thereby provides a principled basis for selecting PAC representations according to the informational demands of the scientific question. The researcher can select coupling strength alone, or choose to interrogate preferred phase, phase polarity, multimodality, and full geometric characterisation. Using analytical derivations, simulations, and empirical EEG data from 2432 subject–channel observations together with an independent EEG dataset from children with DLD and age-matched controls, we show that coupling patterns indistinguishable under conventional metrics become separable within the moment representation. As expected, empirical phase-inversion effects follow the predicted moment transformations. These findings establish PAC as a hierarchy of representations, each preserving different properties of the underlying phase–amplitude structure. This has direct consequences for mechanistic interpretations of cross-frequency coupling in neural systems (Fig. 1a–e).

**Figure 1.**
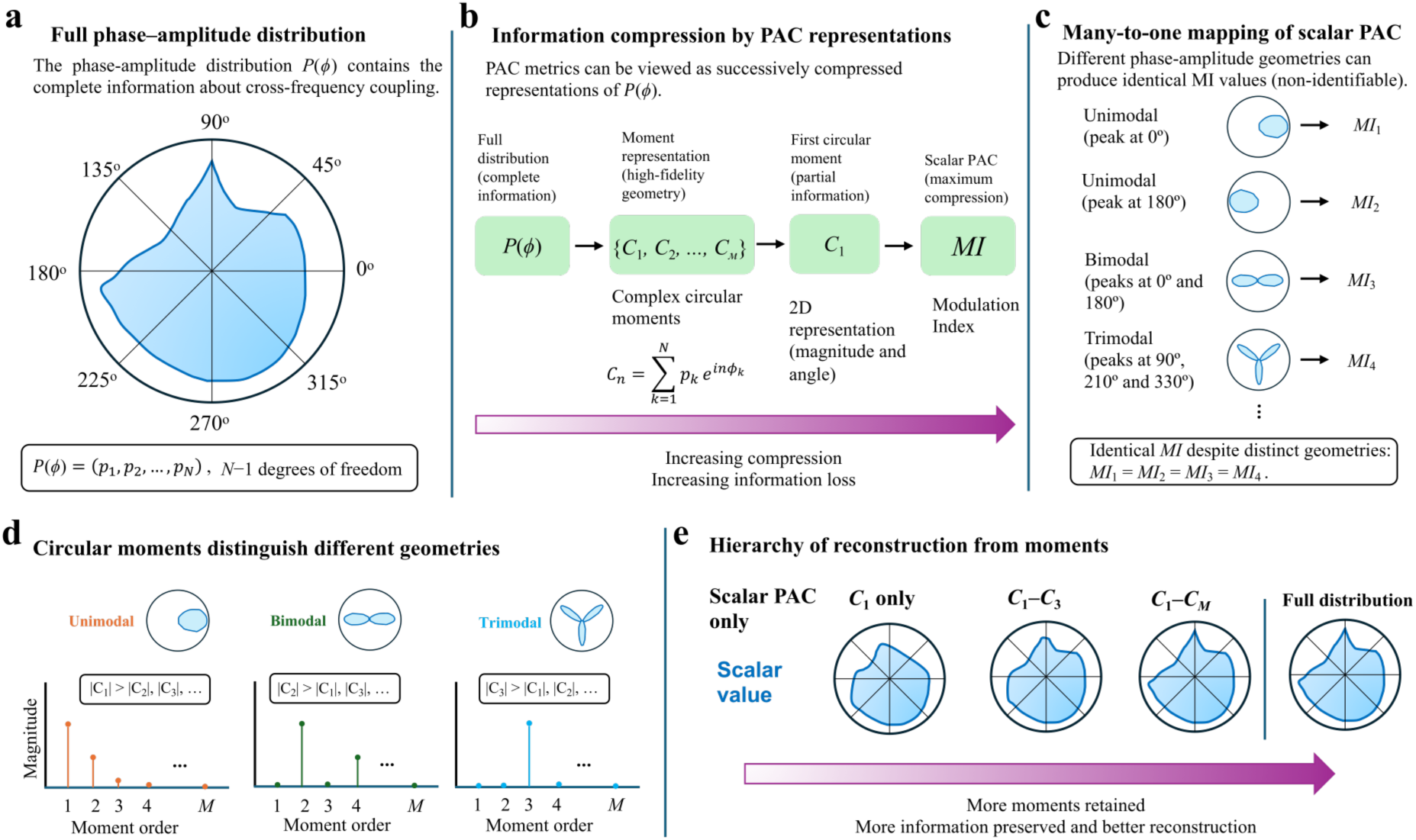
PAC as a hierarchy of information-preserving representations. **(a)** The full phase–amplitude distribution, *P*(*φ*), contains the complete angular structure of cross-frequency coupling and represents the highest-information description. **(b)** Conventional PAC measures can be viewed as compressed representations of *P*(*φ*). Circular moments {*C*_1_, *C*_2_, …, *C_M_*} preserve geometric information, whereas scalar PAC measures such as the MI provide maximally compressed summaries. **(c)** Scalar PAC measures are generally non-identifiable, meaning that distinct phase–amplitude geometries, including unimodal, bimodal, and trimodal structures, can produce identical scalar values. **(d)** Circular moments distinguish these geometries because different coupling structures are expressed at different moment orders. **(e)** Retaining additional circular moments steadily improves reconstruction of the original phase–amplitude distribution, forming a hierarchy between scalar PAC summaries and the full distribution.

## Results

To demonstrate non-identifiability directly, we simulated four phase–amplitude distributions with distinct angular geometries but identical MI values (*MI* = 0.1000; Fig. 2). These comprised a unimodal distribution centred at 0°, a unimodal distribution centred at 180°, a bimodal distribution with peaks at 0° and 180°, and a trimodal distribution with peaks separated by 120°. The concentration parameter of each distribution was adjusted to achieve exact MI matching. Despite identical MI values, the circular moment representation separated the four cases. The two unimodal distributions were dominated by *C*_1_, with equal magnitudes but opposite angular orientations, whereas the bimodal distribution was dominated by *C*_2_ and the trimodal distribution by *C*_3_. These results demonstrate that MI loses phase–amplitude geometry, whereas circular moments preserve structural information that scalar PAC summaries discard.

**Figure 2.**
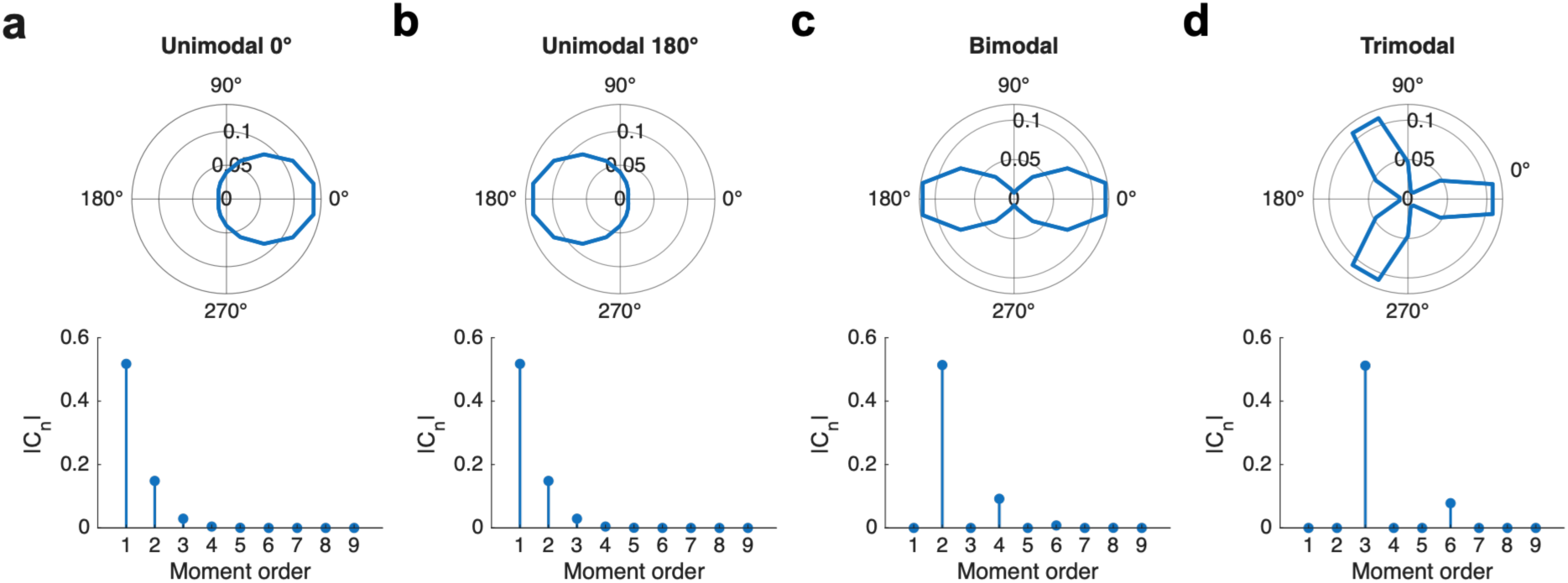
Matched MI values arise from geometrically distinct phase–amplitude distributions. Four simulated phase–amplitude distributions were constructed with different angular structures but identical MI values (*MI* = 0.1000 in all cases). **(a)** Unimodal distribution centred at 0°. **(b)** Unimodal distribution centred at 180°. **(c)** Bimodal distribution with equal peaks at 0° and 180°. **(d)** Trimodal distribution with peaks separated by 120°. For each case, the normalised phase–amplitude distribution is shown as a polar plot and the corresponding circular moment spectrum (|*C_n_*|) is shown as a function of moment order. Despite identical MI values, the four distributions exhibit distinct moment signatures. The unimodal distributions are dominated by *C*_1_, with equal magnitudes but opposite angular orientations, whereas the bimodal and trimodal distributions are dominated by *C*_2_ and *C*_3_, respectively.

### Information recovery increases with moment order

To quantify information retention across the moment hierarchy, we reconstructed a simulated phase–amplitude distribution using successively larger subsets of circular moments (Fig. S1 a–f). Reconstruction based on *C*_1_ alone captured the dominant low-order structure but incompletely recovered finer angular detail (*RMSE* = 0.0082). Including moments *C*_1_–*C*_3_ reduced reconstruction error to *RMSE* = 0.0054, whereas inclusion of moments *C*_1_–*C*_9_ recovered the original 18-bin distribution with negligible error (*RMSE* < 1×10⁻¹⁶). These results demonstrate that circular moments form an information hierarchy in which low-order moments provide compact geometric summaries and higher-order moments recover finer structural detail. Because the phase–amplitude distribution is represented by a finite set of Fourier coefficients, the complete moment set uniquely specifies the original distribution. By contrast, MI compresses the distribution into a single scalar value and therefore does not retain sufficient information to reconstruct its geometry.

### Moment representations reveal information discarded by MI in empirical EEG data

We next examined whether the distinctions predicted by the moment framework are present in empirical electrophysiological recordings obtained during continuous speech listening (DiLiberto et al., 2023). Phase–amplitude distributions were computed and analysed at both the single-observation and population levels. We first investigated representative examples of delta–beta and theta–gamma coupling from channel A10 (Fig. 3a–h). For both frequency pairs, inversion of the low-frequency phase left the MI unchanged (delta–beta: *MI* = 2.71 × 10⁻⁴; theta–gamma: *MI* = 2.25 × 10⁻⁵). The corresponding circular moment representations revealed information that was entirely absent from the scalar PAC estimate. Crucially, moment magnitudes remained unchanged after phase-inversion. In contrast, moment angles transformed systematically according to the theoretical prediction 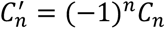, with odd-order moments rotated by approximately 180°. Even-order moments remained unchanged. Although the original and inverted distributions yielded identical MI values, they occupied distinct locations in moment space and exhibited different geometric organisation.

**Figure 3.**
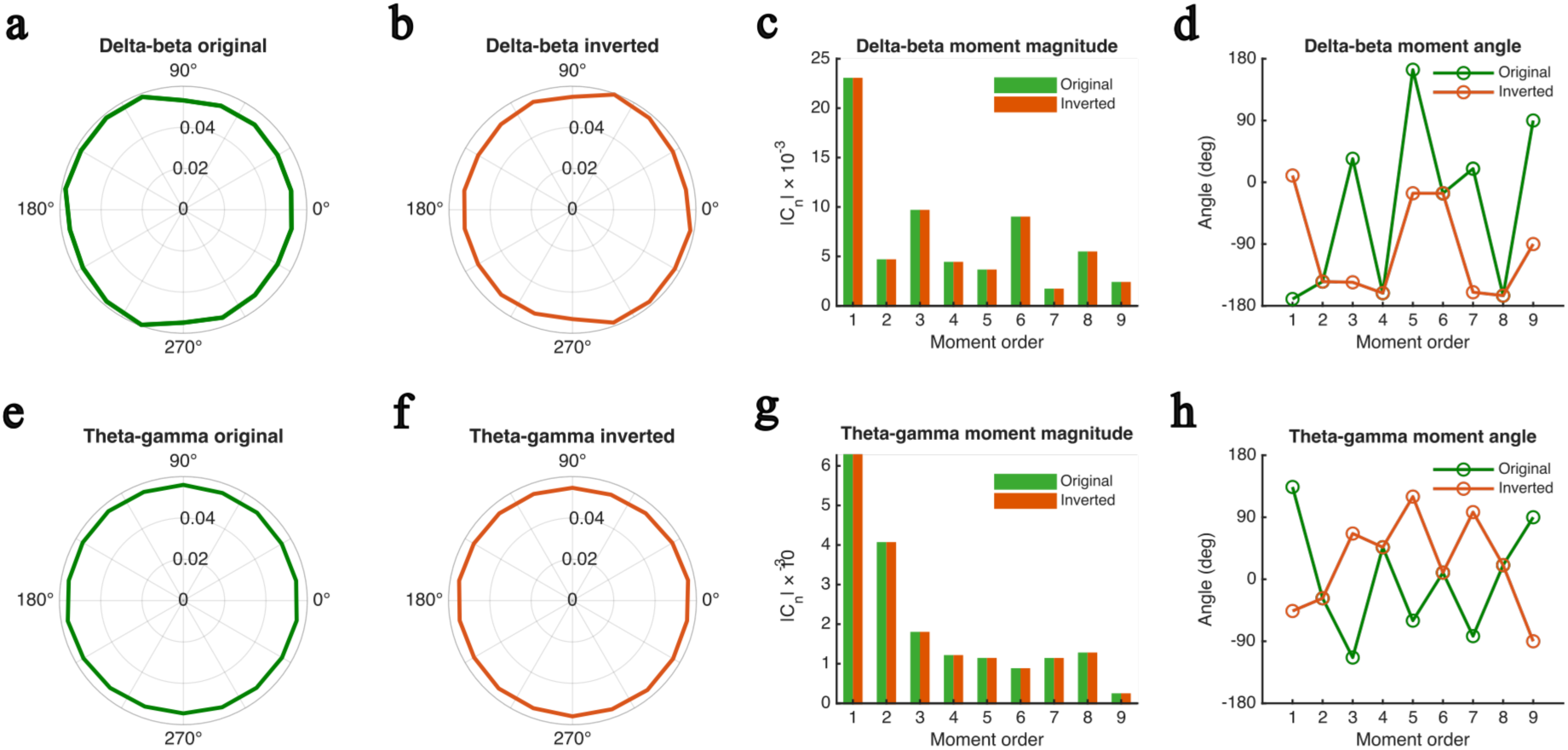
Circular moment representations reveal phase information discarded by MI under phase-inversion in empirical EEG data. Phase–amplitude distributions were computed from a representative EEG channel (A10) during continuous speech listening for **(a–d)** delta–beta and **(e–h)** theta–gamma coupling. **(a, b, e, f)** Polar plots of the normalised phase–amplitude distribution for the original (green) and phase-inverted (orange) low-frequency signals. The two distributions are visually indistinguishable and yield identical MI values (delta–beta: *MI* = 2.71 × 10⁻⁴; theta–gamma: *MI* = 2.25 × 10⁻⁵). **(c, g)** Circular moment magnitudes (|*C_n_*|) for moments 1–9 for the original (green) and inverted (orange) conditions. Moment magnitudes are preserved after phase-inversion, confirming that coupling strength is unaffected. **(d, h)** Moment angles for the original (green) and inverted (orange) conditions. Odd-order moments (*C*_1_, *C*_3_, …) shift by 180° after phase-inversion, whereas even-order moments remain unchanged, in precise agreement with the theoretical transformation 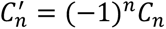.

We next tested whether similar representational ambiguities occur more generally across empirical PAC observations. Analyses were performed on all 2432 subject–channel EEG observations. For both delta–beta (Fig. 4a–e) and theta–gamma coupling (Fig. 4f–j), we identified representative phase–amplitude distributions with nearly identical MI values but markedly different moment structures. In the delta–beta analysis, three representative distributions matched to within 0.7% of a target MI of 1.96 × 10⁻⁴ were dominated by the first-, second-, and third-order moments, respectively. A similar pattern was observed for theta–gamma coupling, where representative distributions matched to within 0.9% of a target MI of 1.48 × 10⁻⁵ exhibited dominant first-, second-, and third-order moment structure despite yielding essentially identical MI values. Accordingly, distinct phase–amplitude geometries can map onto the same scalar PAC estimate.

**Figure 4.**
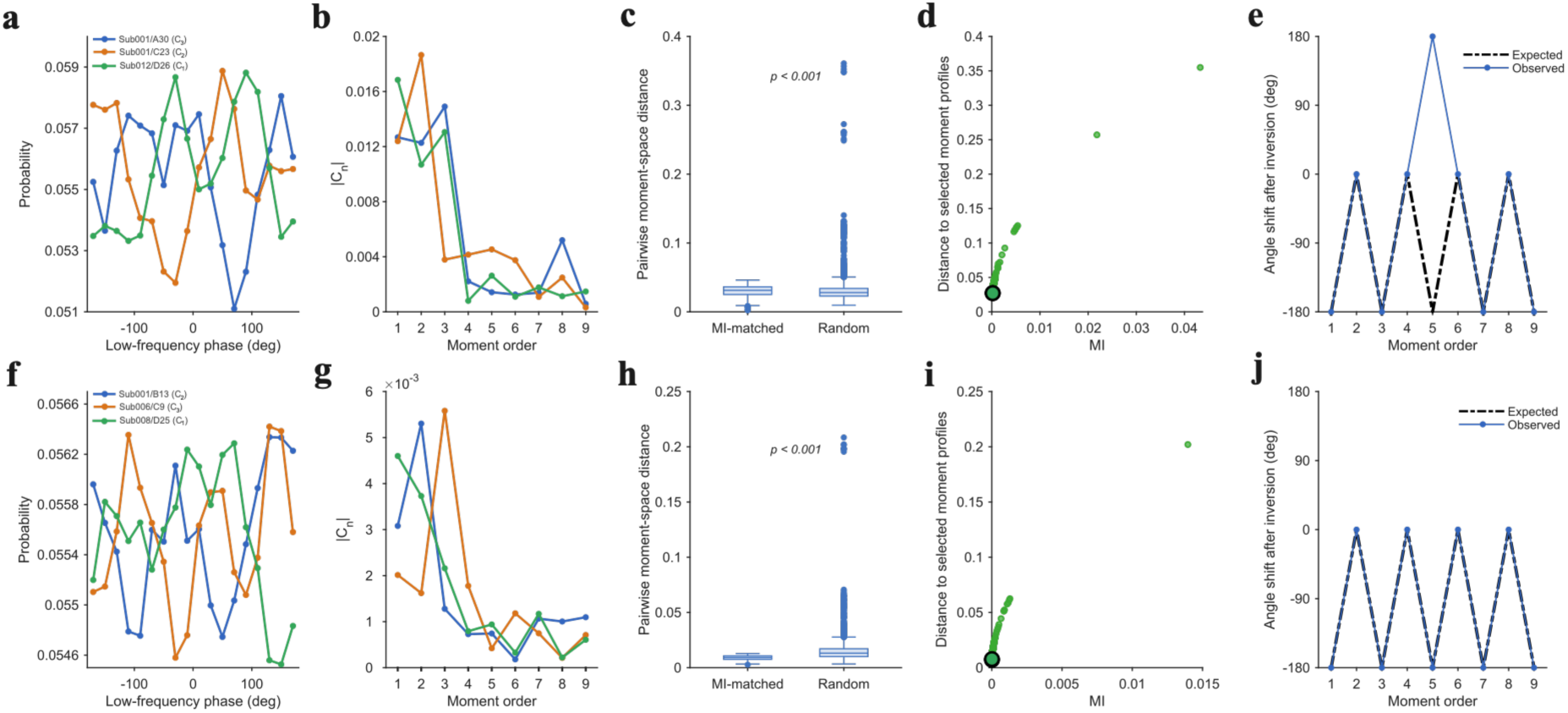
Population-level demonstration of PAC non-identifiability and moment-space geometry for (a–e) delta–beta and (f–j) theta–gamma coupling. **(a,f)** Representative empirical phase–amplitude distributions with nearly identical MI values but different dominant moment structure**. (b, g)** Corresponding moment spectra, showing dominance of the first-, second-, and third-order moments despite comparable MI values. **(c, h)** Distributions of pairwise moment-space distances for MI-matched observations and randomly selected observations. Although constrained to similar PAC strength, MI-matched profiles retain substantial geometric variability. **(d, i)** Moment-space distance as a function of MI across all observations. Profiles with similar MI values occupy widely separated regions of moment space, demonstrating that similarity in PAC strength does not imply similarity in coupling geometry. **(e, j)** Mean angular shift following phase-inversion as a function of moment order. Observed shifts closely follow theoretical predictions, with odd-order moments rotating by 180° and even-order moments remaining unchanged. Across all observations, scalar PAC measures remain invariant while the underlying moment geometry is systematically transformed.

To quantify this effect at the population level, we computed pairwise distances between complete complex-moment representations. Distributions selected within narrow MI windows retained substantial geometric variability despite their nearly identical PAC strength. For delta–beta coupling, the median pairwise moment-space distance among MI-matched observations was 3.13 × 10⁻². For theta–gamma coupling, the corresponding median distance was 9.20 × 10⁻³. Although MI-matched distributions were more similar to one another than randomly selected pairs, the distributions of distances remained broad and statistically distinguishable from the random baseline (two-sample Kolmogorov–Smirnov test, *p* < 0.001 for both frequency pairs; Fig. 4c,h). Furthermore, observations with nearly identical MI values spanned a wide range of moment-space distances (Fig. 4d,i). Clearly, similarity in PAC strength does not imply similarity in phase–amplitude geometry. These results provide population-level evidence that MI constitutes a many-to-one mapping from phase–amplitude distributions to scalar PAC values.

We then evaluated the theoretical prediction that phase-inversion selectively transforms odd- and even-order moments. Across all 2432 subject–channel observations, inversion of the low-frequency phase left MI unchanged, with a maximum absolute difference of only 7.68 × 10⁻¹⁶. In contrast, odd-order moments exhibited the predicted 180° rotation, whereas even-order moments remained unchanged (Fig. 4e,j). The observed angular shifts closely matched theoretical expectations across all moment orders and frequency pairs. Together, these findings demonstrate that scalar PAC measures preserve information about coupling strength while discarding information about phase polarity, multimodality, and temporal organisation. The circular moment representation retains these properties and therefore distinguishes phase–amplitude structures that are indistinguishable under conventional PAC metrics.

To explore whether the moment-based framework can explain the apparently-intact PAC present in children with language disorders, we re-analysed an EEG dataset acquired from children with DLD during continuous speech listening (Keshavarzi et al., 2026). To determine whether comparable coupling strength could conceal differences in the geometry of phase–amplitude coupling, we analysed the first eight complex moments of delta–gamma distribution in a right temporal ROI (Fig. S2 a–d). Delta-gamma PAC was selected on the basis of Keshavarzi et al. (2024), and as opposing auditory theories of DLD predict atypical delta-rate (Goswami, 2022) versus gamma-rate (Tallal, 2004) linguistic processing. MI did not differ between age-matched control and DLD groups (Welch’s t-test, *p* = 0.157). By contrast, two-sample studentised bootstrap analyses of moment direction, with Benjamini–Hochberg correction across *C*_1_ − *C*_8_, identified significant group differences for *C*_1_ and *C*_5_. For *C*_1_, the canonical mean orientation was 312.2° in Controls and 150.81° in DLD, an absolute difference of 161.39° (*p* = 0.010, *q* = 0.041). For *C*_5_, the corresponding orientations were 70.28° and 37.91°, an absolute difference of 32.37° (*p* = 0.004, q = 0.036). No other moment-order comparisons survived correction. These findings provide preliminary evidence that the moment representation can capture group differences in the directional organisation of phase–amplitude distributions that may help explain impaired speech/language processing.

### Statistical robustness and reconstruction properties of moment-based PAC representations

To assess the robustness of the moment-based representation, we examined surrogate inference, bootstrap resampling, data-length convergence, group-level subsampling, and reconstruction performance (Fig. 5a–f). To test whether the dominant moment of a representative empirical PAC profile could be distinguished from chance, a surrogate procedure circularly shifting the high-frequency amplitude envelope relative to the low-frequency phase was applied. The observed first-moment magnitude exceeded the entire surrogate distribution (*p* < 0.001, 1000 surrogates; Fig. 5a), showing that the dominant component of the phase–amplitude distribution was unlikely to arise from chance phase–amplitude associations. Bootstrap resampling revealed a clear hierarchy of moment magnitudes (Fig. 5b). The first moment contributed the largest component of the empirical PAC geometry, and higher-order moments made gradually smaller contributions. The associated 95% confidence intervals were narrow, indicating stable estimation of the moment spectrum across resamples.

**Figure 5.**
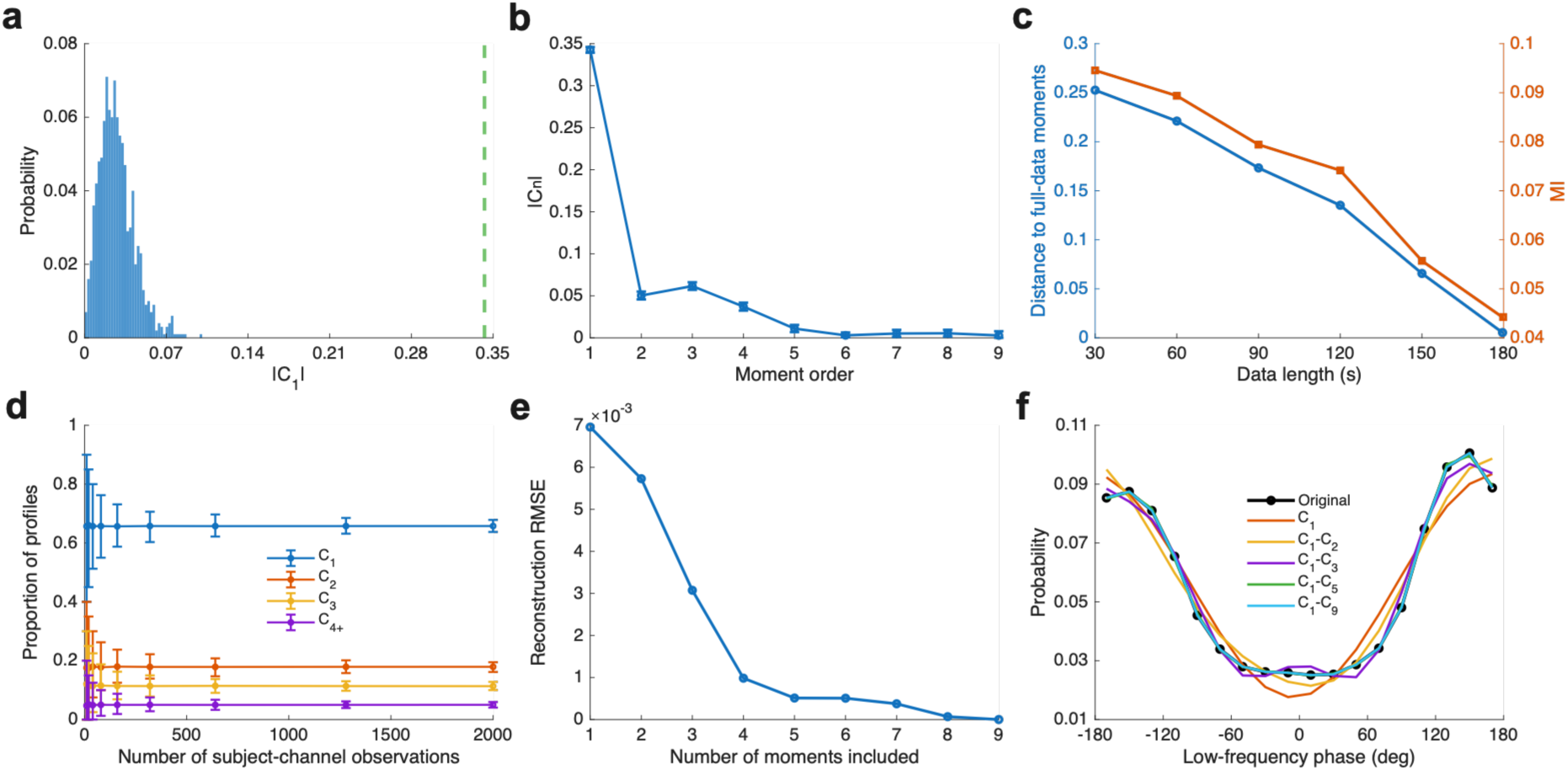
Statistical robustness and reconstruction properties of moment-based PAC representations. Robustness and reconstruction analyses for delta–beta coupling, computed from the empirical EEG dataset. **(a)** Surrogate inference for the dominant first moment of a representative PAC profile. The observed |*C*_1_| (green dashed line) exceeds the surrogate distribution (blue histogram) generated by circularly shifting the high-frequency amplitude envelope relative to the low-frequency phase, indicating a non-random phase–amplitude structure (*p* < 0.001; 1000 surrogates). **(b)** Bootstrap estimates of moment magnitudes (|*C_n_*|) with 95% confidence intervals. The first moment contributes the largest component of the empirical PAC geometry, with diminishing contributions from higher-order moments. **(c)** Data-length convergence. Moment-space distance between representations estimated from longer data segments and the representation estimated from the complete 180-s recording decreases monotonically with increasing data length, while MI gradually approaches the value obtained from the complete recording. **(d)** Dominant-moment composition stability. Subject–channel observations were repeatedly resampled from the full dataset; the estimated proportions of profiles dominated by first- (*C*_1_), second- (*C*_2_), third- (*C*_3_), and higher-order (*C*_4+_) moments converge rapidly and remain stable across increasing sample sizes. Error bars denote 95% bootstrap confidence intervals. **(e)** Reconstruction error as a function of the number of moments included, showing progressive reduction in RMSE as additional moments are retained. **(f)** Moment-based reconstruction of a representative empirical PAC profile. Reconstructions using successively larger moment sets (*C*_1_, *C*_1_–*C*_2_, *C*_1_–*C*_3_, *C*_1_–*C*_5_, and *C*_1_–*C*_9_) recover the original phase–amplitude distribution (black), illustrating how higher-order moments recover finer geometric structure.

We next examined convergence as a function of recording duration. Moment-space distance to the representation estimated from the complete 180-s recording decreased monotonically with increasing data length, while MI steadily approached the value obtained from the complete recording (Fig. 5c). Both measures converged as additional data were incorporated. To assess population-level robustness, we repeatedly resampled subject–channel observations from the full dataset and calculated the proportion of profiles dominated by first-, second-, third-, and higher-order moments. The estimated proportions converged rapidly and remained stable across increasing sample sizes (Fig. 5d). Accordingly, the dominant-moment composition reflected a reproducible property of the dataset and was not driven by a small subset of observations.

Finally, we evaluated reconstruction from finite moment sets. Reconstruction error, quantified using root-mean-square error (RMSE), decreased monotonically as additional moments were included and approached zero when the complete set of moments was retained (Fig. 5e). Correspondingly, reconstructed phase–amplitude profiles more closely approached the original empirical distribution as higher-order moments were added (Fig. 5f). Accordingly, the moment representation is statistically robust, converges with increasing data length, and captures the geometric structure of empirical PAC distributions with increasing fidelity as additional moments are retained.

### A representation-selection framework for future PAC analysis

The preceding analyses establish that PAC can be represented at multiple levels of informational detail, ranging from highly compressed scalar summaries to the complete phase–amplitude distribution. For future studies, we propose a representation-selection framework that formalises PAC as a hierarchy of representations derived from the underlying phase–amplitude distribution (Fig. 6). Within this hierarchy, conventional measures such as MI and MVL correspond to low-dimensional projections that quantify specific properties of the distribution while discarding other aspects of its geometry. By contrast, circular moments preserve richer structural information. The first moment captures preferred phase and phase polarity, higher-order moments encode multimodality and symmetry, and the complete moment representation uniquely specifies the original phase–amplitude distribution. This framework clarifies that no single PAC measure is universally optimal. The appropriate representation instead depends on the scientific question. Scalar measures are suitable when the objective is to quantify coupling strength. Moment-based representations are required when the goal is to characterise phase alignment, multimodality, symmetry, or full coupling geometry. Thus, MI, MVL, preferred phase, and circular moments are not competing measures but complementary representations that occupy different positions along an information-preservation hierarchy.

**Figure 6.**
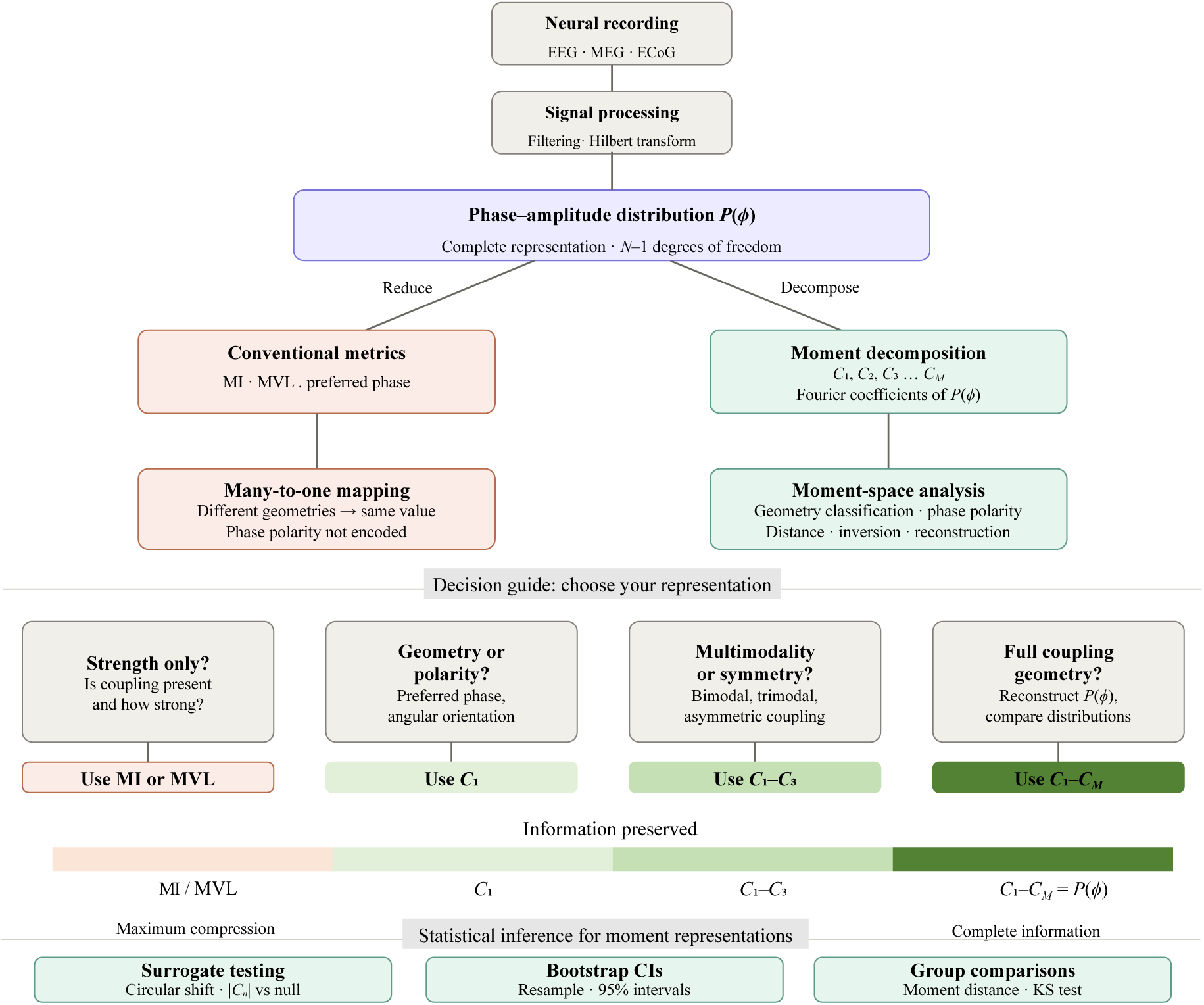
Representation-selection framework for PAC analysis. PAC is conceptualized as a hierarchy of representations derived from the underlying phase–amplitude distribution, *P*(*ϕ*). Conventional PAC measures, including MI, MVL, and preferred phase, correspond to more heavily compressed projections of *P*(*ϕ*) and therefore preserve only selected aspects of the underlying phase–amplitude distribution. Circular moments provide a richer representation in which geometric information is retained across multiple orders: the first moment captures preferred phase and phase polarity, higher-order moments encode multimodality and symmetry, and the complete moment set uniquely recovers the full phase–amplitude distribution. The framework links representation dimensionality to information preservation and provides practical guidance for selecting PAC measures according to the scientific question of interest, ranging from coupling strength and phase polarity to full phase–amplitude geometry.

## Discussion

The present study demonstrates that commonly-used PAC metrics differ fundamentally in the information they preserve about underlying phase–amplitude structure. This is critical when assessing known neural mechanisms of language processing in language-disordered groups. By formalising PAC measures as mappings from phase–amplitude distributions to lower-dimensional representations, we show that information loss is not merely a practical consequence of data reduction but an inherent property of scalar PAC quantification. Conventional measures such as MI and MVL provide compact summaries of phase-dependent amplitude modulation (Canolty et al., 2006; Tort et al., 2010), whereas circular moments retain progressively richer geometric information (Fisher, 1995; Mardia and Jupp, 2000). PAC should therefore be understood as an underlying phase–amplitude distribution that can be represented at different levels of compression.

A key finding is that scalar PAC measures are generally non-identifiable with respect to coupling geometry. Distinct phase–amplitude structures, including unimodal, bimodal, and multimodal profiles, can produce identical MI values despite representing different temporal organisations of neural activity. The phase-inversion invariance of MI, recently established by Keshavarzi (2026) and formalised here within a broader framework, provides a concrete example. Because MI depends on the entropy of the phase–amplitude distribution rather than on the angular arrangement of probability mass, it is insensitive to transformations that preserve the set of bin probabilities while changing their phase ordering.

The circular-moment representation recovers discarded information by expressing the phase–amplitude distribution through its discrete Fourier coefficients. The first moment encodes the preferred phase and distinguishes opposite phase polarities, higher-order moments capture increasingly complex geometric structure, and the complete non-redundant moment set uniquely specifies the original distribution. Reconstruction analyses demonstrated that low-order moments provide compact geometric summaries, whereas additional moments increasingly recover finer angular structure. By contrast, MI provides only a single scalar value and cannot reconstruct the geometry of the underlying distribution.

Empirical analyses of EEG data show that these representational distinctions are present in real neural recordings, with empirical PAC distributions with nearly identical MI values occupying different locations in moment space and exhibiting distinct dominant moment structures. The predicted odd–even transformation of moment phases under phase-inversion was also observed, with odd-order moments rotating by 180° and even-order moments remaining unchanged, while MI remained invariant. These findings validate the theoretical framework and demonstrate that the geometric information captured by circular moments is recoverable from empirical EEG data. The robustness analyses further showed that moment estimates were stable under surrogate testing, bootstrap resampling, increasing recording duration, population subsampling, and moment-based reconstruction.

When applied to an EEG dataset comparing children with DLD and typically-developing controls (Keshavarzi et al., 2026), the circular-moment representation indicated preliminary group differences in the phase geometry of delta–gamma coupling. These differences were not detectable using MI. This insight provides a potential means of reconciling two dominant auditory theories of DLD, one foregrounding delta-rate coupling (Temporal Sampling theory, Goswami, 2022) and one foregrounding gamma-rate speech information (Rapid Auditory Processing theory, Tallal, 2004).

These results have direct implications for all studies in which MI-based PAC strength is interpreted as evidence for temporally structured neural coordination. In oscillatory models of speech processing, PAC plays a central mechanistic role (Giraud and Poeppel, 2012; Hyafil et al., 2015; Arnal et al., 2015; Dogonasheva et al., 2025). For oscillatory models, phase polarity carries mechanistic significance. High-frequency activity peaking at opposite phases of the slow oscillation implies different temporal organisations and different behavioural outcomes. In neural models of working memory and sequential coding, the phase of high-frequency activity relative to the theta cycle is central to accounts of information ordering (Lisman and Jensen, 2013; Daume et al., 2024). However, while MI can quantify the strength of phase-dependent amplitude modulation, it does not encode the phase geometry required to distinguish between temporal organisations.

The findings presented do not invalidate prior MI-based results, rather clarify their scope. The representation-selection framework formalises possible trade-offs. Scalar measures such as MI and MVL remain appropriate for quantifying the presence or strength of phase-dependent modulation. When preferred timing or phase polarity are important, the complex first moment is required. When the research question concerns multimodality or more complex coupling geometry, additional moments become informative. When complete coupling geometries need to be compared or reconstructed, the full moment representation is required. In this framework, MI, MVL, preferred phase, and circular moments are not competing measures but complementary representations of the same underlying phase–amplitude distribution.

Several limitations should be noted. The moment-based framework addresses the representational properties of PAC measures not the validity of PAC estimation itself. Methodological issues such as filtering artefacts, waveform asymmetry, spectral leakage and non-stationarity remain important and should be addressed independently (Aru et al., 2015; Jensen et al., 2016; Cole and Voytek, 2017). Reliable estimation of higher-order moments may require longer recordings than scalar PAC estimates, and the sensitivity of moment estimates to binning resolution warrants further investigation. The number of participants in the DLD dataset was relatively small, so the group-level differences reported should be regarded as requiring replication. Finally, the behaviour of moment representations under low signal-to-noise conditions or strong non-stationarity requires further characterisation.

In conclusion, PAC should be understood as a hierarchy of representations rather than a single scalar metric. The phase–amplitude distribution provides the complete information about coupling geometry, whereas conventional scalar metrics and circular moments preserve different subsets of this information, ranging from phase-dependent modulation strength to phase polarity, multimodality and full geometric structure. By linking representation dimensionality, identifiability, and recoverable information, the suggested framework provides a principled basis for selecting PAC measures according to the temporal properties preserved.

## Methods

### Probabilistic formulation of phase–amplitude coupling

Let *φ*(*t*) ∈ [−*π*, *π*) denote the instantaneous phase of a low-frequency oscillation and let *A*(*t*) ≥ 0 denote the amplitude envelope of a higher-frequency oscillation. The phase interval was partitioned into *N* non-overlapping bins (*B*_1_, *B*_2_, …, *B_N_*) with centres *φ*_1_, *φ*_2_, …, *φ_N_*. For phase bin *B_k_*, the mean amplitude was defined as

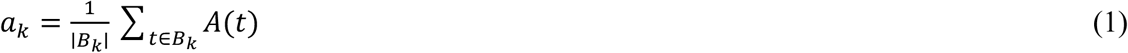

where ∣ *B_k_* ∣ denotes the number of observations assigned to bin *k*. The phase-binned amplitudes were normalised to obtain

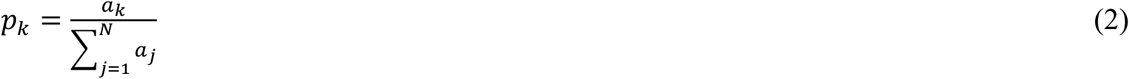

such that 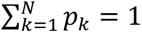. The vector *P* = (*p*_1_, *p*_2_, …, *p_N_*) therefore defines a discrete probability distribution over low-frequency phase. Throughout this work, *P* is treated as the fundamental representation of phase–amplitude coupling. All PAC measures considered in this study are derived as mappings from *P* to lower-dimensional representations.

### PAC measures as representational mappings

We formalise a PAC measure as a mapping

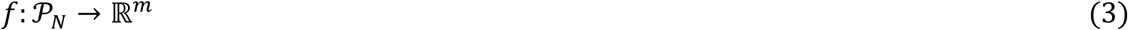

where *P_N_* denotes the space of discrete phase–amplitude distributions over *N* phase bins. Given a distribution *P*, the measure produces the representation

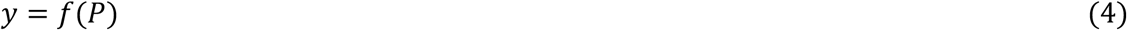

Different PAC measures correspond to different mappings from the same underlying phase–amplitude distribution and may therefore preserve different amounts of information about the original structure. For conventional scalar PAC metrics, *m* = 1, so the phase–amplitude distribution is compressed into a single value. The same formulation also includes vector-valued representations, such as the circular moment hierarchy, for which *m* > 1. A representation is identifiable if

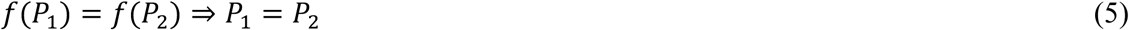

Conversely, if there exist distinct distributions *P*_1_ ≠ *P*_2_ such that *f*(*P*_1_) = *f*(*P*_2_), then the representation is non-identifiable and discards information about the underlying phase–amplitude structure.

### Modulation index

Following Tort et al. (2010), the modulation index (MI) was defined as

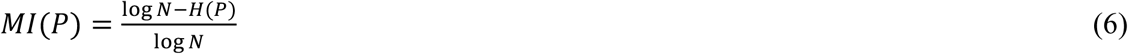

where *H*(*P*) is the Shannon entropy of the phase–amplitude distribution,

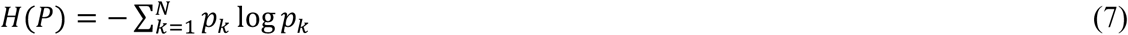

MI quantifies the deviation of *P* from the uniform distribution and is commonly interpreted as a measure of PAC strength. Within the present representational framework, MI maps the full phase–amplitude distribution to a single scalar value. Consequently, all information retained by MI is mediated through the entropy of *P*.

Entropy depends on the set of probability values {*p*_1_, *p*_2_, …, *p_N_*}, not on their angular arrangement across phase bins. Therefore, distributions with different phase organisations but identical entropy yield identical MI values. MI thus preserves information about the degree of phase-dependent modulation while discarding information about phase geometry. This property makes MI generally non-identifiable. Distinct unimodal, bimodal, or multimodal phase–amplitude distributions can produce identical MI values when their entropies are matched. MI therefore quantifies deviation from uniformity but does not uniquely specify the angular geometry or temporal organisation of phase–amplitude coupling.

##### Circular moment representation

**Proposition 1 (Identifiability of the complete moment representation).** Let *N* ≥ 2 be an integer, and let *P* = (*p*_1_, …, *p_N_*) be a real-valued phase–amplitude distribution discretised into *N* equally spaced phase bins with centres

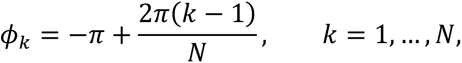

satisfying

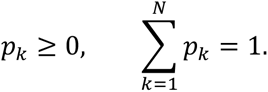

Define the circular moments as

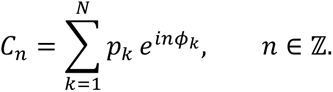

Then the non-redundant set of moments

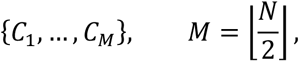

uniquely determines *P*. The negative-order moments are obtained from the conjugate-symmetry relation

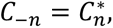

where (⋅)^∗^ denotes complex conjugation and *C*_0_ = 1. Although the moments are defined for all integer orders, only the finite non-redundant set specified above is required for reconstruction. For odd *N*, the corresponding inverse reconstruction is

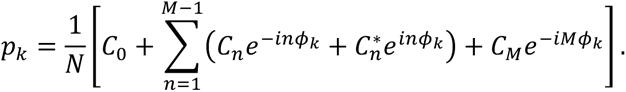

For even *N*, the inverse reconstruction is

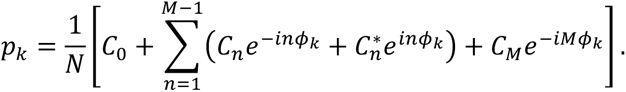

When *N* is even, *C_M_* = *C_N_*_/2_ is the real-valued Nyquist coefficient and is included only once.

Because the circular moments are the discrete Fourier coefficients of *P*, the invertibility of the discrete Fourier transform guarantees that the complete non-redundant moment set uniquely specifies the original phase–amplitude distribution. The complete moment representation is therefore injective, whereas scalar PAC measures such as MI and MVL are generally non-identifiable because distinct distributions can produce identical metric values. The first circular moment, *C*_1_, captures the dominant angular structure of the phase–amplitude distribution. Its magnitude, |*C*_1_|, quantifies concentration around a preferred phase, whereas its argument, arg(*C*_1_), specifies angular orientation. Higher-order moments, *C_n_* for *n* > 1, capture increasingly complex geometric properties, including multimodality, symmetry, and higher-order structure. The first circular moment is closely related to the mean resultant vector underlying MVL, but retains both magnitude and angular orientation. This distinction is important because *C*_1_ vanishes for distributions symmetric under a 180° phase shift, making MVL insensitive to symmetric bimodal coupling patterns that may still yield non-zero MI values. Conversely, MI is invariant to preferred phase and phase polarity and therefore discards the angular information encoded by *arg*(*C*_1_).

### Information content and reconstruction

For a phase–amplitude distribution discretized into *N* phase bins, the normalized probability vector *P* contains *N*–1 independent degrees of freedom because 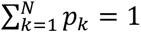. Since the circular moments are equivalent to the Fourier coefficients of the distribution, a sufficiently large set of moments uniquely specifies *P*. In practice, however, low-order moments capture the dominant geometric features of the phase–amplitude structure. The first moment characterises preferred phase and concentration, the second moment captures bimodal and symmetric structure, and higher-order moments encode finer-scale features of the distribution. Consequently, the number of moments required depends on the desired reconstruction fidelity and the complexity of the underlying coupling pattern.

### Transformation under phase rotation

Consider a rotation of the phase axis by angle *α*, such that 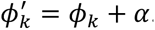. The corresponding circular moment is

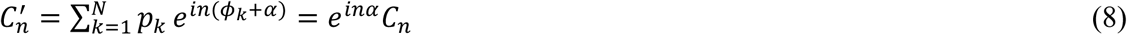

Thus, circular moments transform predictably under phase rotations. For phase-inversion, *α* = *π*, this gives

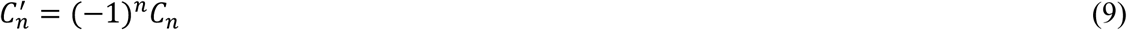

Odd-order moments therefore change sign, whereas even-order moments remain unchanged. In particular, 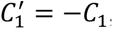, so the complex first moment encodes phase polarity, whereas its magnitude |*C*_1_|, MVL, and MI remain invariant. Scalar PAC summaries therefore preserve coupling strength but do not encode the phase polarity retained by the complex moment representation.

### Empirical EEG dataset

Empirical validation analyses were performed using a publicly available EEG dataset collected during continuous speech listening (Di Liberto et al., 2023). EEG was recorded from 19 participants using a 64-channel BioSemi ActiveTwo system at a sampling rate of 512 Hz. Data preprocessing followed the procedures described in the original publication. Low-frequency phase signals were extracted from the delta (1–4 Hz) and theta (4–8 Hz) bands, and high-frequency amplitude envelopes were extracted from the beta (13–25 Hz) and gamma (25–40 Hz) bands using band-pass filtering and the Hilbert transform. PAC representations were computed independently for all participant–channel combinations for both frequency pairs (delta–beta and theta–gamma), yielding 2432 observations (19 participants × 64 channels × 2 frequency pairs). These data were used to evaluate the representational properties of PAC measures in empirical electrophysiological recordings.

Generalisability was further examined using an independent preprocessed EEG dataset acquired during continuous speech listening from 32 children, comprising 16 children with developmental language disorder (DLD) and 16 age-matched typically developing controls (Keshavarzi et al., 2026). Analyses were conducted at a sampling rate of 100 Hz within a predefined 12-channel right temporal ROI (channels 100, 102, 109, 114, 121, 116, 101, 107, 108, 115, 113, and 120) used by Keshavarzi et al. (2026). For each channel, mean low-gamma amplitude was calculated within 18 equally spaced delta-phase bins and normalised to form a phase–amplitude distribution. Channel-level MI values were averaged to obtain one participant-level MI estimate, whereas the normalised phase–amplitude distributions were first averaged across valid ROI channels and then used to calculate participant-level moments *C*_1_–*C*_8_. Group differences in MI were assessed using a two-sided Welch’s independent-samples *t*-test. Moment directions were compared using two-sided two-sample studentised bootstrap tests with 10000 resamples, followed by Benjamini–Hochberg correction across *C*_1_–*C*_8_. Statistical testing was performed on the full circular direction arg(*C_k_*), and division by *k* was used only to express canonical orientation differences in the original phase domain.

### Statistical analyses

Statistical robustness was evaluated using surrogate testing, bootstrap resampling, and subsampling analyses. Surrogate distributions were generated by circularly shifting the high-frequency amplitude envelope relative to the low-frequency phase for 1000 iterations. Bootstrap confidence intervals were estimated using 1000 resamples. Differences between distributions of moment-space distances were assessed using two-sided two-sample Kolmogorov–Smirnov tests. Moment-space distance between two phase–amplitude distributions, *P_i_* and *P*_j_, was defined as the Euclidean distance between their complete complex-moment vectors,

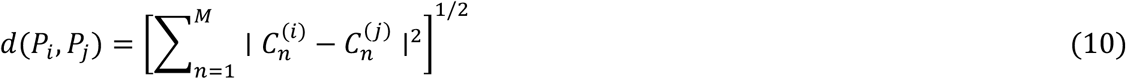

Distances were computed using unnormalised complex-moment vectors and therefore reflected differences in both moment magnitude and moment phase. Matched-set distances were defined as pairwise distances among observations falling within the selected MI window, whereas the random baseline comprised pairwise distances between observations sampled across the full MI range. Reconstruction accuracy was quantified using root-mean-square error (RMSE) between the original and reconstructed phase–amplitude distributions.

## Author contributions

M.K. conceived the study, developed the theoretical framework, performed the mathematical analyses, implemented the computational methods, conducted the simulations and empirical analyses, generated the figures, interpreted the results, and wrote the manuscript. U.G. contributed to conceptualisation and to writing of the manuscript.

## Supplementary Information

**Figure S1.**
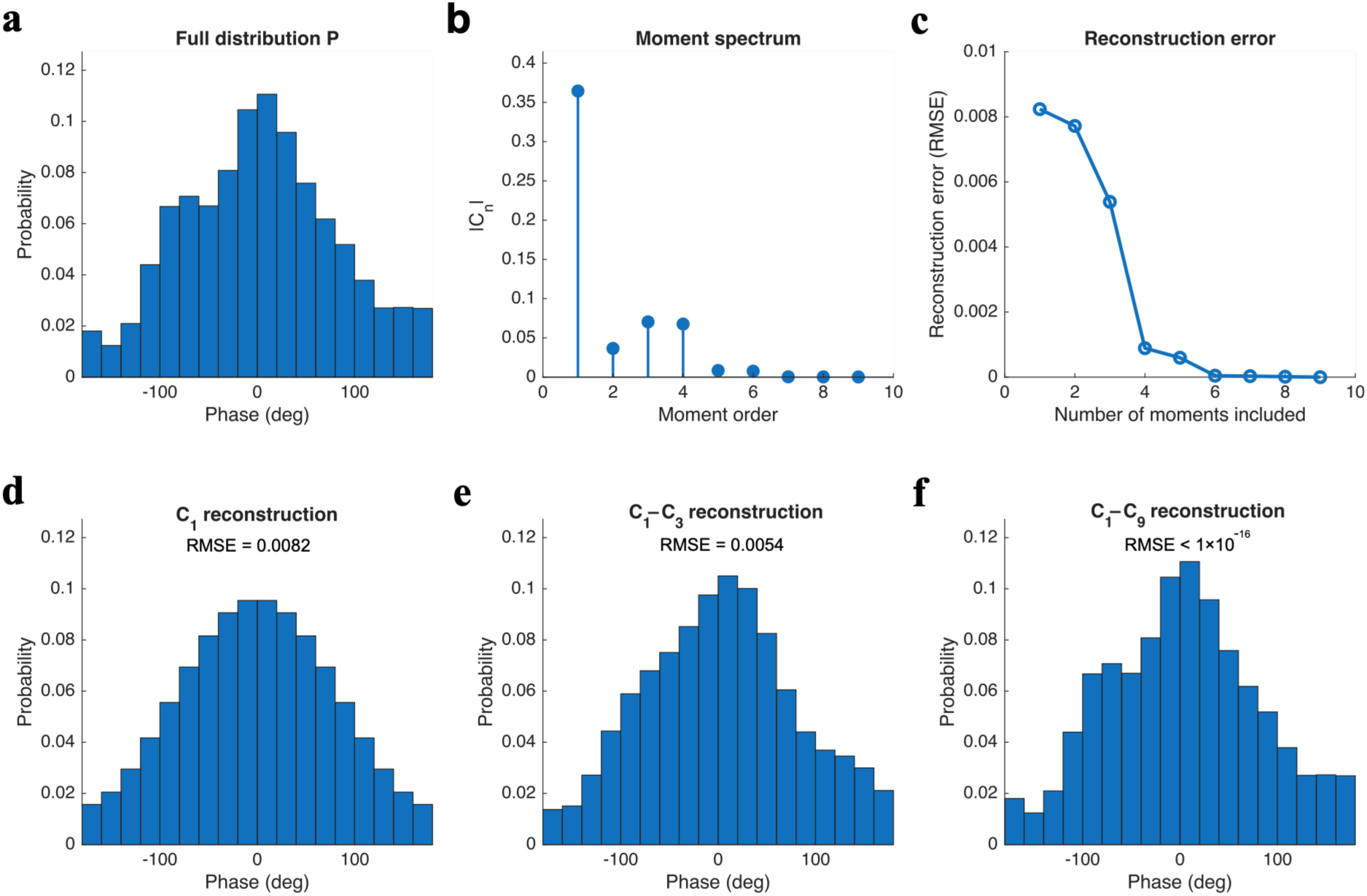
Circular moments form a hierarchy of information recovery. **(a)** Simulated phase–amplitude distribution, *P*(*φ*), comprising the complete 18-bin angular representation. **(b)** Circular moment spectrum (|*C_n_*|), showing that the dominant contribution arises from *C*_1_, with progressively smaller contributions from higher-order moments. **(c)** Reconstruction error as a function of the number of moments retained, decreasing monotonically as additional moments are included. **(d)** Reconstruction using *C*_1_ alone (*RMSE* = 0.0082). **(e)** Reconstruction using *C*_1_–*C*_3_ (*RMSE* = 0.0054). **(f)** Reconstruction using *C*_1_–*C*_9_ (*RMSE* < 1 × 10⁻¹⁶).

**Figure S2.**
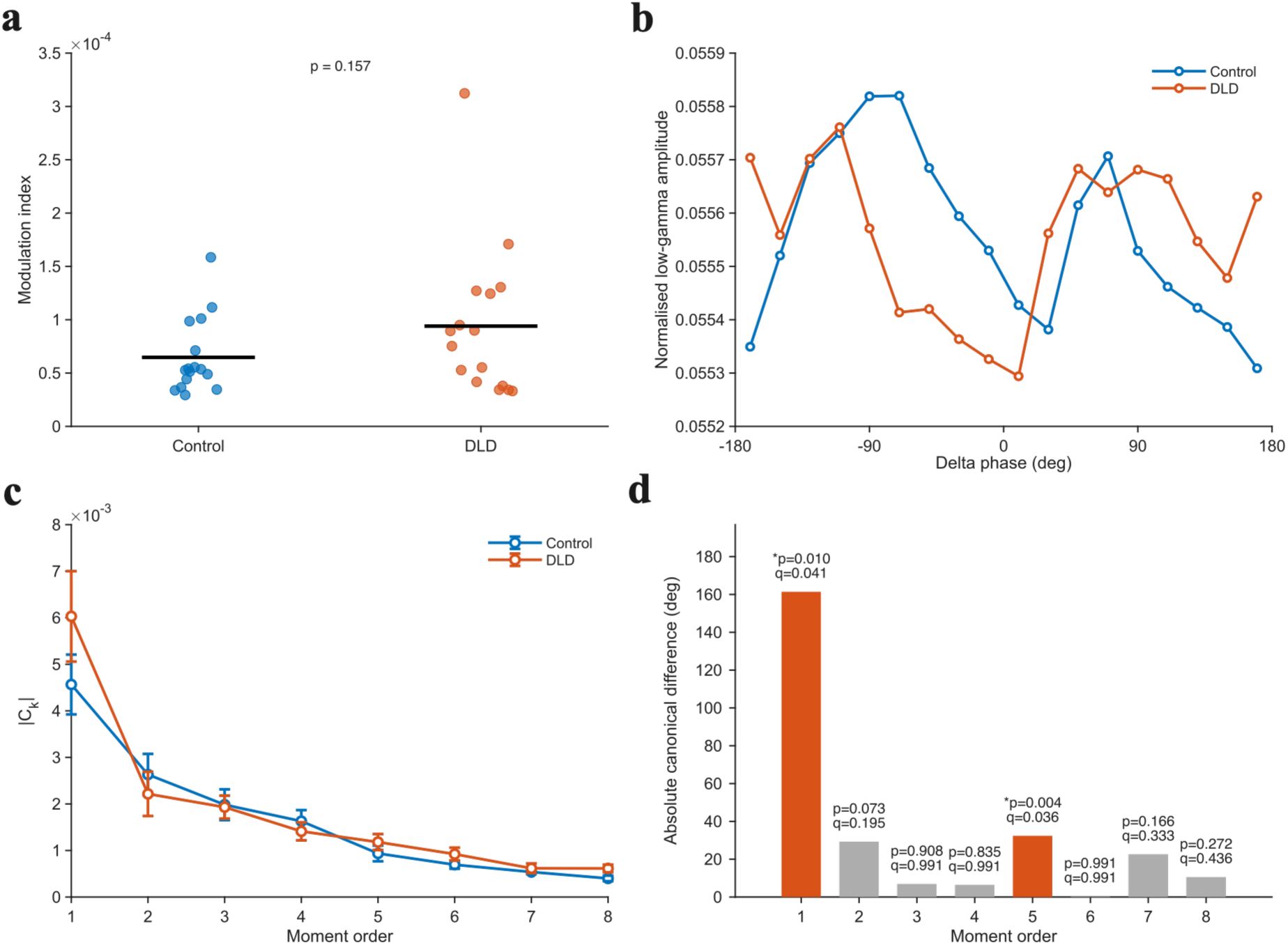
Moment-based analysis of delta–gamma distribution in the right temporal region of interest. Phase–amplitude distributions were calculated using delta phase and low-gamma amplitude from children with developmental language disorder (DLD; *n* = 16) and age-matched typically developing controls (*n* = 16). **(a)** Participant-level modulation index (MI) values. Dots represent individual participants and horizontal black lines indicate group means. MI did not differ significantly between groups. **(b)** Group-average normalised gamma amplitude as a function of delta phase. **(c)** Mean magnitudes of complex moments *C*_1_–*C*_8_. Error bars indicate the standard error of the mean across participants. **(d)** Absolute canonical orientation differences between groups. Statistical testing was performed on the circular mean directions of *C_k_* in moment space. The resulting shortest angular differences were divided by *k* only for representation in the original phase domain. Group differences in moment angle direction were assessed using two-sided two-sample studentised bootstrap tests with 10000 resamples. Labels show raw *p* values and Benjamini–Hochberg-adjusted *q* values across *C*_1_–*C*_8_, orange bars and asterisks denote comparisons surviving FDR correction at *q* < 0.05. Significant differences were observed for *C*_1_ and *C*_5_.

